# Patterned photocatalytic proximity labelling for spatial interactomics

**DOI:** 10.64898/2026.06.01.729422

**Authors:** Takeharu Mino, Hideki Nakamura, Tomonori Tamura, Itaru Hamachi

## Abstract

Mapping protein–protein interactions and proximity within their native spatial context is essential for elucidating molecular mechanisms underlying biological processes; however, methods for spatial interactomics with subanatomical resolution are limited. Here, we introduce digital micromirror device (DMD)–PhoxID, a method that combines photocatalytic proximity labelling with patterned irradiation using a DMD to map the proximal molecular environment of proteins of interest within defined subregions of fixed brain tissue.

## Main

Protein–protein interaction (PPI) networks differ across major brain regions, such as the cortex, hippocampus and cerebellum.^1-3^ Furthermore, expression variation across anatomically and functionally specialized subregions suggests differences in local interaction networks.^4,5^ However, differences in PPIs across subregions have not been characterized. This is mainly because conventional methods for PPI studies involve disruption of brain tissue architecture. Recently developed enzyme-based proximity labelling (PL) enables PPI analysis while preserving the cellular connectivity of brain tissue; however, substrate diffusion limits precise spatial control.^3,6^ Although photocatalyst-mediated PL holds promise for spatial interactome analysis, a dedicated workflow has not been established.^7^

Herein, we report a platform for protein interactome mapping in fixed brain tissues at subregional resolution. The method integrates PhoxID, a genetic manipulation-free photocatalytic PL,^3^ with spatially controllable photoirradiation using a digital micromirror device (DMD) (Fig. 1a). In the DMD–PhoxID protocol, the photosensitizer monobromofluorescein (MBF) is covalently attached to a protein of interest (POI) using a ligand-directed probe.^8^ Fixed brain slices are subjected to spatially confined light irradiation in the presence of a nucleophilic reagent to label proteins proximal to the POI. Labelled proteins can be extracted using SDS-containing high-pH buffer and identified by conventional LC–MSMS. Importantly, this workflow is compatible with immunostained tissue slices, allowing DMD irradiation focused on visually distinguishable brain regions and subregions defined by specific molecular markers.

**Fig. 1.**
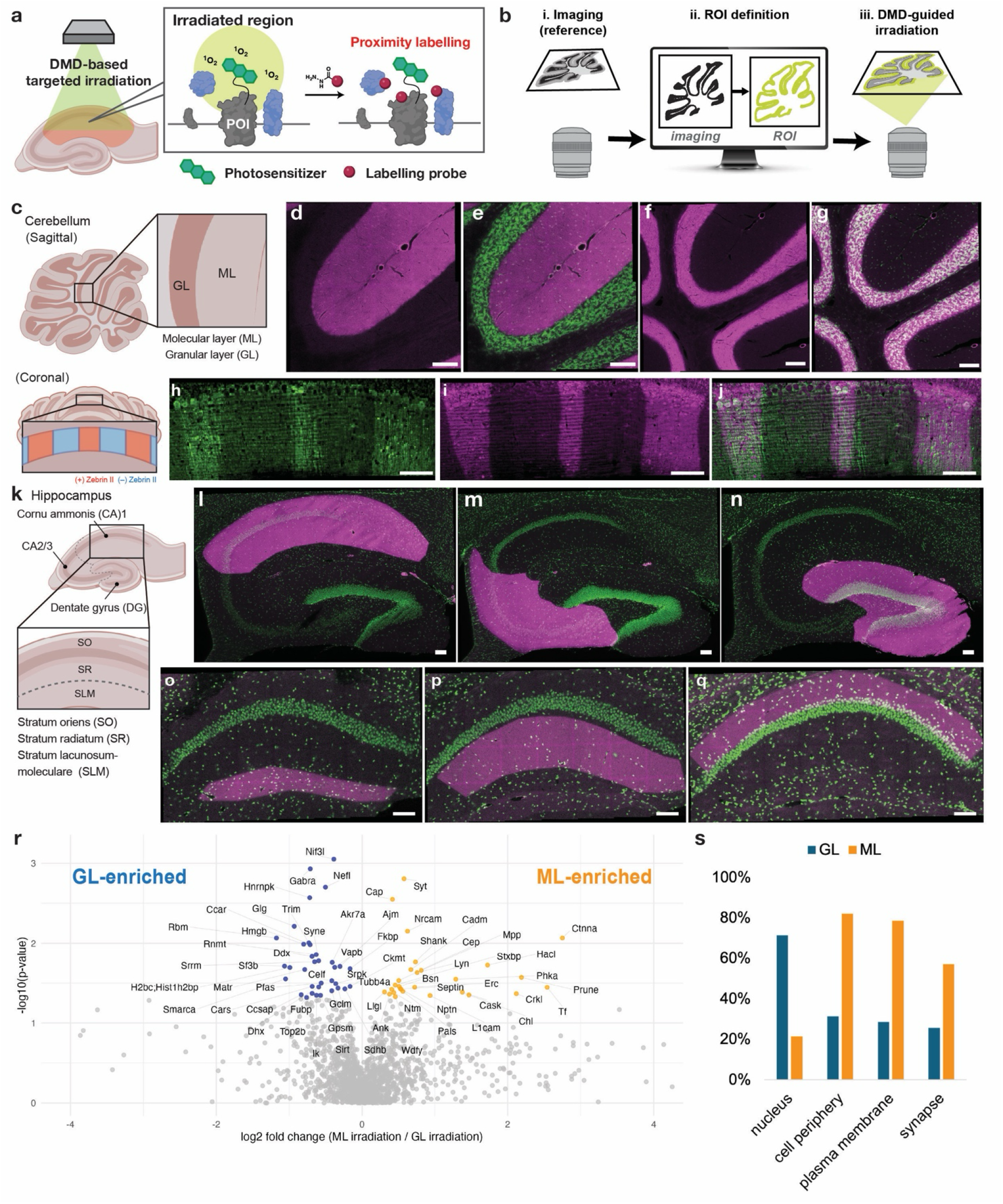
DMD-guided irradiation enables spatially confined proximity labelling in brain tissue. **a**, Concept of DMD–PhoxID. A photosensitizer anchored to proteins generates singlet oxygen upon patterned irradiation, enabling proximity labelling within defined regions. **b**, Workflow for spatial targeting. Reference imaging (i) is used to define regions of interest (ROIs) (ii), followed by DMD-guided irradiation (iii). **c, k**, Structure of the cerebellum (c), hippocampus (k) and their subregions. **d–g**, Confocal images of cerebellar slices following irradiation. Ax647–hydrazide labelling (magenta) is confined to the irradiated ML (d,e) or GL (f,g), with nuclear staining in green. **h**–**j**, Fluorescence imaging of cerebellar slices labelled with Ax647 hydrazide (magenta) following DMD-guided irradiation with regions defined by Zebrin immunostaining (green). **l–q**, Region- and layer-specific labelling in hippocampal slices following irradiation of CA1, CA2/3, DG, SLM, SR and SO.d–j, l–q, Scale bar: 100 µm. **r**, Volcano plot of proteins enriched in ML- and GL-irradiated samples. **s**, Subcellular localization of enriched proteins based on Gene Ontology cellular component annotation.

To evaluate the spatial resolution of DMD–PhoxID, paraformaldehyde-fixed mouse cerebellar slices were modified with MBF–NHS, followed by nuclear staining and bright-field imaging. The molecular layer (ML, nuclear staining–negative) and granule cell layer (GL, nuclear staining–positive) were delineated using an image-based automatic segmentation pipeline (Fig. 1b, c). After DMD irradiation in the presence of a fluorescent nucleophilic probe, AlexaFluor647–hydrazide, light-induced protein labelling was confined to the irradiated ML or GL (Fig. 1d–g, Extended Data Fig. 1, 2). Application to the Zebrin stripe in the ML demonstrated that labelling can be induced at the subanatomic level with single-cell resolution (Fig. 1h–j, Extended Data Fig. 3).^9^ Similar spatial labelling was achieved in the hippocampal cornu Ammonis 1 (CA1), CA2/3 and dentate gyrus (DG) as well as within CA1 subfields, including the stratum lacunosum-moleculare (SLM), stratum radiatum (SR) and stratum oriens (SO) (Fig. 1k–q).

ML-specific irradiation in the presence of biotin–hydrazide enriched synaptic and membrane-associated proteins, whereas GL irradiation preferentially recovered nuclear and RNA-binding proteins (Fig. 1r, s, Supplementary Table 1). These data are consistent with the neuromorphological organization in which the ML is primarily composed of granule cell parallel fibres and Purkinje cell dendrites, whereas the GL is densely populated by granule cell somata, supporting the ability to resolve the proteomic composition of subanatomical brain regions.

To verify the ability of DMD–PhoxID to capture proteins proximal to a POI, we targeted α-amino-3-hydroxy-5-methyl-4-isoxazolepropionic acid-type glutamate receptor (AMPAR), which is distributed across hippocampal regions and localized to excitatory postsynapses (Extended Data Fig. 4). Endogenous AMPAR was selectively modified with MBF in the live mouse brain, followed by fixation, sectioning, and subfield-specific irradiation in hippocampal CA1, CA2/3 and DG, with four biological replicates. Many enriched proteins were annotated to excitatory synapses, demonstrating AMPAR-centred PL (Fig. 2a–d). Heatmaps revealed subfield-dependent patterns (Fig. 2e). Several AMPAR-associated proteins, including Dlg4, Cacng8 and Olfm1, showed identical fold-change (FC) values across all subfields. In contrast, Nptx1 and Plxna1 were enriched in CA1 and CA2/3 but reduced in DG, consistent with known expression patterns and with proximity ligation assay results (Fig. 2h–l; Extended Data Fig. 5, Supplementary Table 2).

**Fig. 2.**
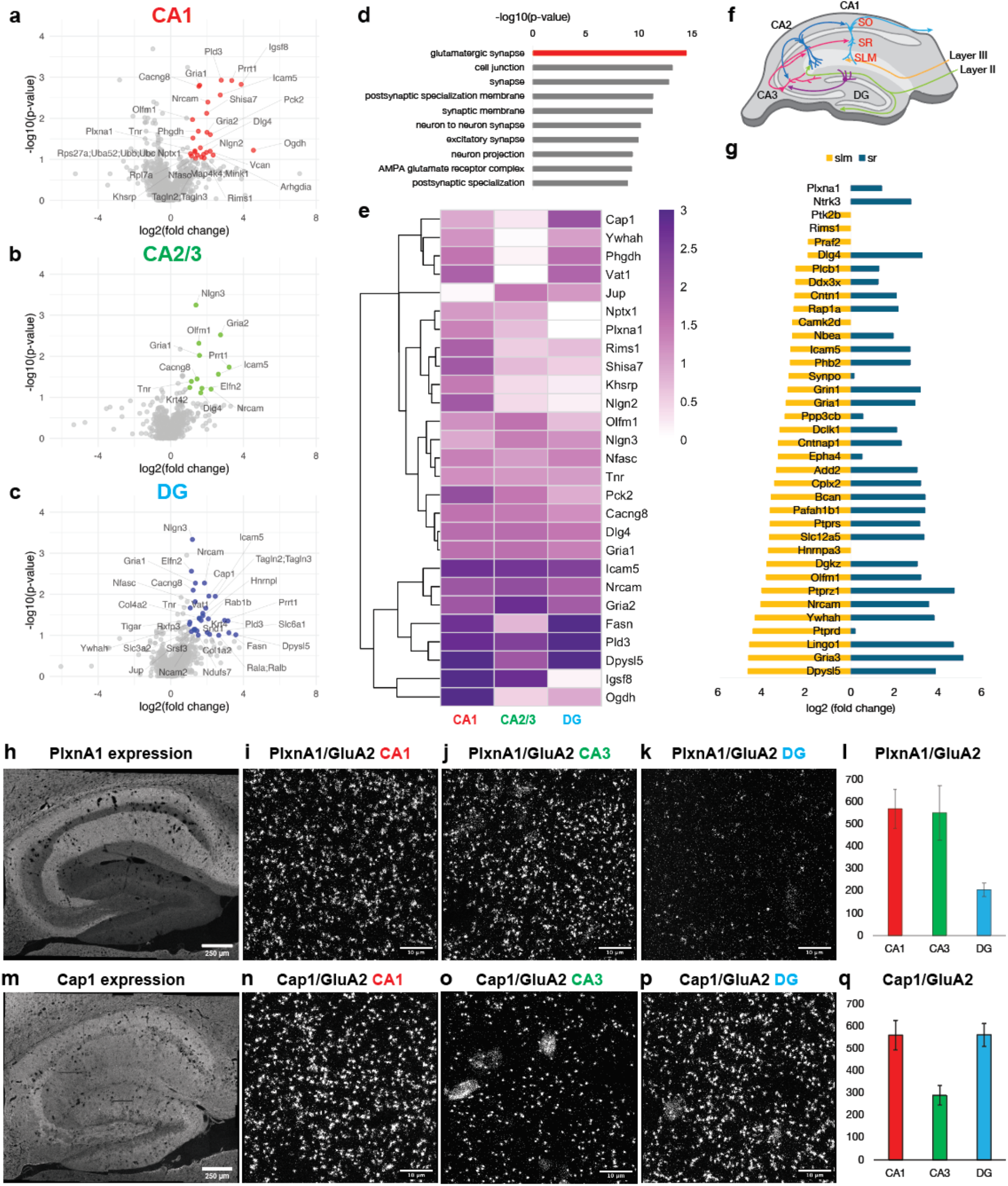
Subfield- and layer-resolved AMPAR-proximal environments revealed by DMD–PhoxID. **a–c**, Volcano plots of proteins enriched in CA1 (a), CA2/3 (b) and DG (c) relative to non-irradiated controls (log_2_(fold-change) versus –log_10_(p-value)). **d**, Gene Ontology cellular component analysis of enriched proteins. **e**, Heatmap of glutamatergic synapse-associated proteins across subfields. **f**, Schematic of hippocampal circuitry and laminar organization. **g**, Differential enrichment of AMPAR-proximal proteins between SLM and SR, revealing layer-specific molecular environments within CA1. **h**,**m**, Immunofluorescence of Plxna1 (h) and Cap1 (m). **i–k, n–p**, PLA images of Plxna1–GluA2 (i–k) and Cap1–GluA2 (n–p) in CA1, CA3 and DG. **l**,**q**, Quantification of PLA puncta. Data are means ± s.e.m.

Notably, although adenylyl cyclase-associated protein 1 (Cap1) is uniformly expressed throughout the hippocampus, the DMD–PhoxID FC was markedly reduced in CA2/3 (Fig. 2e, m, Supplementary Table 2). We hypothesized that this discrepancy arises from subregion-specific differences in Cap1–AMPAR proximity. In support of this, proximity ligation assays revealed reduced Cap1–GluA2 (an AMPAR subunit) proximity in CA3 compared with CA1 and DG (Fig. 2n–q). Given the role of Cap1 in actin regulation and postsynaptic structural plasticity, its reduced proximity in CA3 is consistent with predominant presynaptic plasticity in this subfield.^10^

We performed DMD–PhoxID in SLM and SR of CA1, which are difficult to separate anatomically (Fig. 2f, g). Ntrk3 and Plxna1 were SR-specific, consistent with previous results.^11^ Nine proteins were identified as SLM-specific (FC in SR < 1.2), including Ptprd and Praf2, known to be more abundant in SL.^12,13^ Synaptopodin (Synpo), identified as SLM-selective using DMD–PhoxID, has been reported to exhibit similar expression levels in SLM and SR, despite more Synpo-positive spines proximal to AMPAR in SLM.^14^ These data demonstrate that DMD–PhoxID can capture proximity differences that cannot be predicted from protein abundance distributions.

In summary, DMD–PhoxID enables localized PL and quantitative profiling of subregion-dependent protein proximity in the brain. The established workflow will be flexibly applied to various proteins and non-brain tissue sections using different ligands in the photosensitizer-targeting probe or antibodies. DMD–PhoxID provides a complementary dimension to existing spatial transcriptomics and spatial proteomics approaches^15,16^ for analyses of molecular organization in anatomical microdomains.

## Acknowledgements

The authors would like to thank M. Ishikawa for his support in chemical synthesis. We thank Edanz (https://jp.edanz.com/ac) for editing a draft of this manuscript. This work was supported by the Japan Society for the Promotion of Science Grant-in-Aid for Specially Promoted Research 23H05405 to I.H.; the JST FOREST program (JPMJFR230G) to T.T.; the JST PRESTO programs (JPMJPR24OD and JPMJPR21EB), and the Japan Society for the Promotion of Science Grant-in Aid for Scientific Research (B) (general) (JP26247756) to H. N.; the Japan Society for the Promotion of Science Grant-in-Aid for JSPS Fellows (23KJ1289) to T.M..

## Author contributions

T.M., H.N., T.T., and I.H. conceived the project and designed the experiments. T.M. performed the experiments and data analysis. T.M., T.T., and I.H. wrote the manuscript with input from all authors.

## Methods

### General materials and methods for biochemical/biological experiments

Unless otherwise noted, all reagents were obtained from commercial suppliers (Tokyo Chemical Industry (TCI) or Thermo Fisher Scientific) and used without further purification. SDS-PAGE and western blotting were carried out using a Bio-Rad Mini-PROTEAN III electrophoresis apparatus. Samples were applied to SDS-PAGE and electrotransferred onto polyvinylidene fluoride membranes (Bio-Rad), followed by blocking with 5% nonfat dry milk in Tris-buffered saline containing 0.05% Tween 20 (TBS-T). Biotinylated proteins were detected using streptavidin-HRP conjugate (Invitrogen, S911, 1:4000). Chemiluminescent signals generated with ECL Prime (Cytiva) were detected using a Fusion Solo S imaging system (Vilber Lourmat).

### General information for fluorescence imaging experiments

Confocal laser scanning microscopy (CLSM) was performed using a TCS SP8 confocal microscope (Leica Microsystems, Germany) equipped with a 10× objective lens (numerical aperture (NA) = 0.40, dry objective) and a 100× objective lens (NA = 1.40, oil objective), together with a GaAsP detector. Excitation light was derived from a white laser and set to an appropriate wavelength depending on the fluorophore.

### Animals

Five-week-old male C57BL/6N mice were purchased from Japan SLC, Inc. (Shizuoka, Japan) and maintained under specific pathogen-free conditions. Animals were housed in a controlled environment (23 ± 1 °C, 12 h light/dark cycle) with ad libitum access to food and water, in accordance with the Guidance for Proper Conduct of Animal Experiments issued by the Ministry of Education, Culture, Sports, Science and Technology of Japan. All experimental procedures were performed in accordance with the National Institutes of Health Guide for the Care and Use of Laboratory Animals and were approved by the Institutional Animal Care and Use Committees of Kyoto University.

### Anchoring of photosensitisers in the mouse brain

C57BL/6N mice (male, 5 to 7 weeks old, body weight 18 to 25 g) were anaesthetised by intraperitoneal injection of a mixture of Domitor® (Nippon Zenyaku Kogyo Co., Ltd.), midazolam (Sandoz) and Vetorphale® (Meiji Seika Pharma Co., Ltd.). CAM2–MBF^3^ solution (4.4 µL, 100 µM in PBS(−) containing 5% DMSO) was directly injected into the left and right lateral ventricles (anteroposterior (AP), −1.3 mm from bregma; mediolateral (ML), ±2.0 mm; depth, 2.0 mm) using a microinjector (Nanoliter 2010, World Precision Instruments) at a rate of 600 nL min^−1^ through burr holes drilled in the skull. After injection, the capillary was left in place for 1 min and then slowly withdrawn. The scalp was sutured, and the anaesthetic antagonist Antisedan® (Nippon Zenyaku Kogyo Co., Ltd., 0.2 mL) was administered by intraperitoneal injection. Mice were returned to their home cages and maintained for 13 h to allow covalent modification of endogenous AMPA-type glutamate receptors (AMPARs) by CAM2–MBF.

### Perfusion fixation and brain section preparation

C57BL/6N mice (male, 5 weeks old, body weight 18 to 23 g) were deeply anaesthetised with isoflurane and transcardially perfused with ice-cold 4% paraformaldehyde (PFA) in PBS(−) (pH 7.4, 60 mL). After dissection, brains were further fixed in 4% PFA/PBS(−) overnight at 4 °C. For photo-proximity labelling experiments, fixed brains were washed three times with PBS(−) and sectioned into sagittal cerebellar or hippocampal slices (100 or 250 µm thick) using a vibratome (LinearSlicer Pro7, Dosaka EM). For immunostaining and proximity ligation assay experiments, fixed brains were washed with PBS(−), sagittally divided along the midline, and immersed in 30% sucrose in PBS(−) at 4 °C for 1 to 3 days. Brain tissues were embedded in optimal cutting temperature (OCT) compound, frozen, and sectioned sagittally at a thickness of 15 µm using a cryostat (CM1950, Leica Biosystems).

### Photo-proximity labelling for fluorescence imaging

Sagittal cerebellar or hippocampal slices (100 or 250 µm thick) were incubated in a solution containing Hoechst 33258 (10 µg mL^−1^) and MBF–NHS (100 µM) in PBS(−) for 2 h at room temperature in a 48-well plate (Falcon) to achieve non-targeted protein modification. After incubation, slices were washed three times with PBS(−) and immersed in Alexa Fluor 647 hydrazide (10 µM, Invitrogen) in PBS(−) (75 µL) for 5 to 20 min at room temperature in 35 mm glass-bottom dishes (Matsunami, whole-glass type). Slices were subsequently covered with glass coverslips (Matsunami, 18 mm × 18 mm, No.1), and fluorescence imaging was performed using an inverted fluorescence microscope (Eclipse Ti2, Nikon) equipped with a 4× dry objective lens (NA = 0.20). Fluorescence signals were collected using a CMOS sensor (Hamamatsu, C14440-20UP). Excitation was provided by an LED light source (X-Cite, Excelitas) with an ET365/10x excitation filter (Chroma) for Hoechst 33258 imaging. Regions of interest (ROIs) were manually or semi-automatically defined on the basis of Hoechst 33258 fluorescence images or immunostaining signals. Irradiation patterns were generated in situ using NIS-Elements software and projected onto the tissue using a DMD-based irradiation system. Patterned photoirradiation was performed using a multipattern LED irradiation system (Optoline, Leopard2, LEO2-K-RI-NK-UHPI460) coupled to a digital micromirror device (DMD). Defined ROIs were selectively illuminated to induce local photooxidation. After photoirradiation, slices were washed three times with PBS(−). Slices were further washed overnight with 0.1% Triton X-100 in PBS(−) (5 mL) to remove unreacted reagents before imaging by confocal laser scanning microscopy. Confocal images were acquired under identical settings within each experiment for quantitative comparison.

### Photo-proximity labelling for proteomic analysis

Coronal hippocampal slices prepared from brains of CAM2–MBF-treated mice were incubated with Hoechst 33258 (10 µg mL^−1^) in PBS(−) for 30 min at room temperature. Slices were subsequently immersed in hydrazide–PEG4–biotin (Hyd–PEG4–Bt, 10 mM) in PBS(−) (75 µL) in 35 mm glass-bottom dishes (Matsunami, whole-glass type) and covered with glass coverslips (18 mm × 18 mm, No.1). ROIs were defined on the basis of Hoechst 33258 fluorescence images, and patterned photoirradiation was performed using the same DMD-based irradiation system described above. After photoirradiation, slices were washed three times with PBS(−) and homogenized in 500 mM Tris-HCl buffer (pH 8.5) containing 6% SDS using an ultrasonic cell disruptor (Branson Ultrasonics, Sonifier SFX 250). The lysates were incubated at 95 °C for 2 h, mixed with 10 volumes of chilled acetone (proteomics grade), and stored at −20 °C overnight. Protein precipitates were collected by centrifugation (15,000 × g, 10 min, 4 °C), washed with an additional 10 volumes of acetone by centrifugation (3,000 × g, 1 min, 4 °C), and air-dried at room temperature for 10 min. Precipitates were dissolved in 300 μL of 1% SDS RIPA buffer (pH 7.4, 25 mM Tris-HCl, 150 mM NaCl, 1% SDS, 1% Nonidet P-40, and 0.25% deoxycholic acid) by sonication using an ultrasonic cell disruptor. The solution was diluted with an equal volume of 0.1% SDS RIPA buffer (pH 7.4, 25 mM Tris-HCl, 150 mM NaCl, 0.1% SDS, 1% Nonidet P-40, and 0.25% deoxycholic acid) to reduce the SDS concentration to approximately 0.5%, further sonicated, and heated at 45 °C for 45 min. Biotinylated proteins were subsequently enriched using High Capacity NeutrAvidin Agarose beads as described below.

### Enrichment of labelled proteins

Following centrifugation of the lysates (17,730g, 10 min, 4 °C), the protein concentration of the supernatant was determined using a BCA assay. Samples were diluted to a final protein concentration of 1.5 mg mL^−1^ using approximately 0.5% SDS-containing RIPA buffer. A total of 0.7 to 2 mg protein was incubated with High Capacity NeutrAvidin Agarose beads (100 µL bed volume) pre-washed with PBS(−) in 1.5 mL Protein LoBind tubes (Eppendorf). After rotation at room temperature for 2 h, beads were collected by centrifugation (3,000g, 1 min, 4 °C) and washed four times with RIPA buffer. Beads were transferred to fresh 1.5 mL Protein LoBind tubes and washed an additional four times with RIPA buffer. For elution, beads were heated with 25 µL of 5× sample buffer containing 250 mM DTT at 95 °C for 5 min. Beads were removed using Micro Bio-Spin columns (Bio-Rad), and the eluates were collected. Labelled proteins were detected by western blotting using streptavidin-HRP conjugate (Invitrogen, S911, 1:4000).

### Sample preparation for NanoLC-MS/MS

Enriched protein samples were loaded onto 10% SDS–PAGE gels (Mini-PROTEAN TGX Gels, Bio-Rad) and electrophoresed for approximately 1 cm. Gel regions containing proteins were manually excised, diced, and fixed in 45% methanol/water containing 5% acetic acid for 20 min. Fixed gel pieces were washed sequentially with 50% aqueous methanol and ultrapure water, followed by dehydration with acetonitrile. Dehydrated gels were rehydrated with 200 µL of 10 mM DTT in 100 mM triethylammonium bicarbonate (TEAB) buffer (pH 8.5) and incubated at 56 °C for 30 min. The DTT solution was replaced with 55 mM iodoacetamide in 100 mM TEAB buffer, and samples were incubated at 37 °C in the dark for 30 min. Gels were subsequently dehydrated with acetonitrile, rehydrated with 100 mM TEAB buffer, and dehydrated again with acetonitrile. Gel pieces were then swollen in 100 mM TEAB buffer containing Sequence Grade Trypsin (Promega, V5113; 10 ng µL^−1^) and incubated overnight at 37 °C for in-gel digestion. Peptides were extracted by addition of 50% acetonitrile containing 0.1% trifluoroacetic acid (TFA), followed by sonication for 10 min in a bath sonicator. The supernatant was collected into fresh Protein LoBind tubes. This extraction step was repeated once with 50% acetonitrile containing 0.1% TFA and twice with 100% acetonitrile containing 0.1% TFA. Combined peptide extracts were concentrated using a centrifugal concentrator (CC-105, TOMY) and further purified using GL-Tip SDB columns (GL Sciences) according to the manufacturer’s instructions.

### NanoLC-MS/MS analysis

NanoLC-MS/MS analyses in data-independent acquisition (DIA) mode were performed using an Orbitrap Exploris 480 mass spectrometer (Thermo Fisher Scientific) coupled to a Vanquish Neo UHPLC system (Thermo Fisher Scientific). Peptides were loaded onto a nano-HPLC capillary column (0.075 × 125 mm, packed with 3 µm C18 particles; Nikkyo Technos, Tokyo, Japan) via a PepMap Neo trap cartridge (Thermo Fisher Scientific). The injection volume was 2 µL and the flow rate was maintained at 300 nL min^−1^. Mobile phases consisted of (A) 0.5% acetic acid in water and (B) 0.5% acetic acid in 80% acetonitrile. Peptides were separated using a multistep linear gradient of 5 to 5% B over 3.6 min, 5 to 19% B over 40.7 min, 19 to 29% B over 15.5 min, 29 to 45% B over 10.2 min, 45 to 95% B over 2 min, followed by 95% B for 18 min. A spray voltage of 1.6 kV, ion transfer tube temperature of 275 °C, and normalised HCD collision energy of 30% were applied. The precursor mass scan range and isolation window were set to m/z 500 to 860 and 6 m/z, respectively. Orbitrap resolution was set to 30,000. Raw MS data were analysed using DIA-NN version 1.8.1. In silico spectral libraries were generated from FASTA files using deep learning-based spectral and retention time prediction against the Mus musculus UniProt/Swiss-Prot database (release 2025_02_06). Trypsin/P specificity with up to one missed cleavage was allowed. Peptide lengths of 5 to 30 amino acids were permitted. The precursor charge range was set to 1 to 4, precursor m/z range to 500 to 860, and fragment ion m/z range to 200 to 1,500. Match-between-runs (MBR) was not implemented, whereas isotopologue detection was enabled. Precursor false discovery rate (FDR) was controlled at 1%. Carbamidomethylation of cysteine residues was specified as a fixed modification, and N-terminal methionine excision was allowed as a variable modification. Protein grouping was performed using heuristic protein inference to minimise redundancy and improve consistency across datasets. Two technical LC-MS/MS replicates were acquired for each biological sample. Missing values were imputed using random values sampled from the lowest 5% of detected intensities in each raw file. Geometric means of technical replicates were calculated to obtain protein abundance values for each biological replicate (n = 3 or 4). Statistical analyses and data visualisation were performed in R using custom scripts.

### Gene Ontology and subcellular localisation analysis

Gene Ontology (GO) cellular component enrichment analysis was performed on region-enriched protein sets using the detected proteome as the background dataset. Subcellular localisation annotations were obtained from UniProt and related public databases.

### Immunostaining fixed brain slices

Fixed brain slices prepared as described above were mounted on glass slides and air-dried for at least 30 min. Heat-induced epitope retrieval was performed by autoclaving sections in citrate buffer (0.01 M, pH 6.0) at 120 °C for 20 min. Sections were permeabilised with PBS containing 0.1% Triton X-100 for 15 min at room temperature, washed with PBS, and blocked with PBS containing 0.1% Triton X-100, 10% normal donkey serum (NDS), and 10% normal goat serum (NGS) for 30 min at room temperature. Primary antibodies were diluted in PBS containing 0.1% Triton X-100 and incubated overnight at 4 °C. The following primary antibodies were used: mouse anti-GluA2 (Merck Millipore, MAB397, clone 6C4, 1:500), rabbit anti-TARPγ8 (Frontier Institute, MSFR105820, 1:100), goat anti-Plxna1 (R&D Systems, AF4309-SP, 1:200), and rabbit anti-CAP1 (Abcam, ab96354, 1:500). After PBS washes, sections were incubated for 1 h at room temperature with Alexa Fluor-conjugated secondary antibodies, including goat anti-mouse IgG Alexa Fluor 555 (Abcam, ab150114, 1:200), goat anti-rabbit IgG Alexa Fluor 647 (Invitrogen, A-21422, 1:200), and donkey anti-goat IgG Alexa Fluor 647 (Abcam, ab150131, 1:200). Sections were rinsed briefly in Milli-Q water, air-dried, mounted using Fluoromount PLUS mounting medium, cover-slipped, and imaged by confocal laser scanning microscopy.

### Proximity ligation assay

Protein proximity was assessed using the Duolink® in situ proximity ligation assay (PLA; Sigma-Aldrich) according to the manufacturer’s instructions. Fixed brain slices prepared as described above were mounted on glass slides and air-dried for at least 30 min. Heat-induced epitope retrieval was performed by autoclaving sections in citrate buffer (0.01 M, pH 6.0) at 120 °C for 20 min. Sections were permeabilised with PBS containing 0.1% Triton X-100 for 15 min at room temperature and washed with PBS. Tissue sections were enclosed using a hydrophobic barrier pen and blocked with Duolink® blocking solution for 1 h at 37 °C. Primary antibodies were diluted in PBS containing 0.1% Triton X-100 and incubated overnight at 4 °C. The following antibody combinations were used for PLA analysis: mouse anti-GluA2 (Merck Millipore, MAB397, clone 6C4, 1:500) with rabbit anti-CAP1 (Abcam, ab96354, 1:500), mouse anti-GluA2 with goat anti-Plxna1 (R&D Systems, AF4309-SP, 1:200), and mouse anti-GluA2 with rabbit anti-TARPγ8 (Frontier Institute, MSFR105820, 1:100). After washing with Duolink® wash buffer A, Duolink® PLA probes anti-rabbit PLUS and anti-mouse MINUS were applied according to the manufacturer’s instructions. For goat primary antibodies, anti-goat PLUS PLA probes were used in combination with anti-mouse MINUS probes. Ligation and rolling-circle amplification reactions were subsequently performed using the supplied reagents. Amplified PLA signals were detected using Texas Red-labelled oligonucleotides. Sections were rinsed briefly in Milli-Q water, air-dried, mounted using Fluoromount PLUS mounting medium, cover-slipped, and imaged by confocal laser scanning microscopy. Negative control samples lacking one primary antibody were included to assess background signal.

### Quantification and statistical analysis

Fluorescence intensity and PLA puncta were quantified using Fiji/ImageJ. Biological replicates were defined as independent animals. Statistical analyses were performed using two-sided unpaired Welch’s t-tests unless otherwise indicated. Data are presented as mean ± s.e.m.

## Data availability

The mass spectrometry proteomics data generated in this study, including raw files and DIA-NN analysis outputs, have been deposited to the ProteomeXchange Consortium (https://proteomecentral.proteomexchange.org) via the jPOST partner repository (https://jpostdb.org/) with dataset identifiers JPST004636 and PXD078507^17^.

All data supporting the findings of this study are available within the paper and its Supplementary Information. Source data are provided with this paper.

## Extended Data Figures

**Extended Data Fig. 1:**
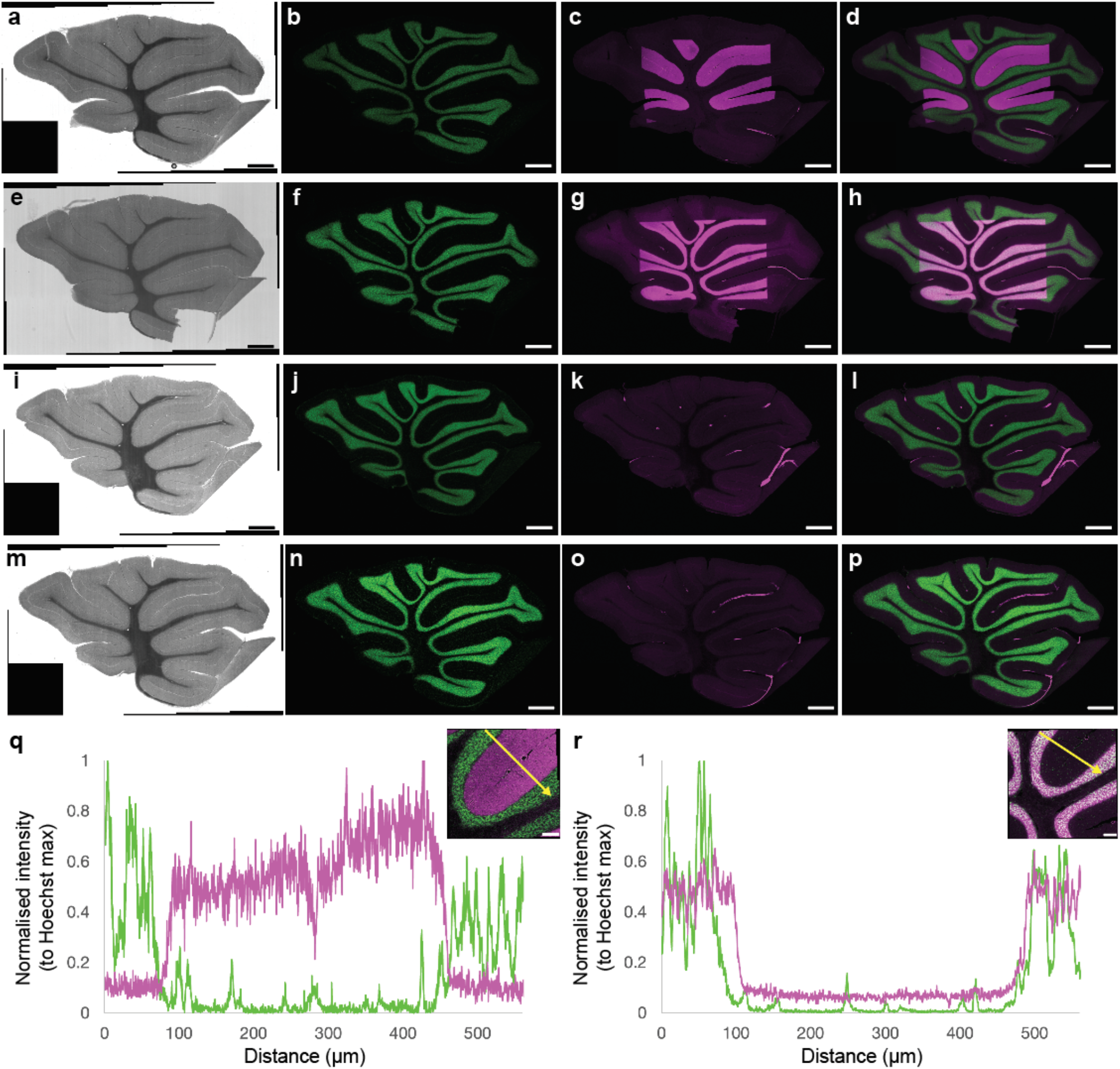
Spatially confined MBF-mediated labelling in cerebellar slices and control conditions. **a–d**, Fluorescence imaging of cerebellar slices following DMD-guided irradiation of the molecular layer (ML) in the presence of MBF–NHS, corresponding to Fig. 1d,e. **e–h**, Fluorescence imaging following irradiation of the granule cell layer (GL) under the same conditions, corresponding to Fig. 1f,g. **i–l**, Control experiment without MBF–NHS, showing minimal labelling upon ML irradiation. **m–p**, Control experiment with MBF–NHS but without light irradiation, showing negligible labelling. In all panels, Ax647–hydrazide labelling is shown in magenta, nuclear staining in green, and differential interference contrast (DIC) images are shown in greyscale. These results demonstrate that labelling is dependent on both MBF and light irradiation and is spatially confined to the irradiated regions. PFA-fixed cerebellar slices were incubated with MBF–NHS (100 µM, 2 h, room temperature) and irradiated in PBS containing Ax647–hydrazide (10 µM) using DMD-patterned 520 nm light for 30 min. After irradiation, samples were washed overnight in 0.1% Triton X-100 in PBS and imaged by confocal microscopy. **q**,**r**, Line profile analysis for samples irradiated in ML (q) or GL (r), corresponding to Fig. 1d,e and Fig. 1f,g, respectively. Intensity profiles were obtained along lines traversing ML and GL, as indicated by yellow arrows in the insets, and normalised to the maximum Hoechst signal within each line. Mean profiles from n = 8 lines per region. These profiles confirm that MBF-mediated labelling is spatially confined to the irradiated layer. Scale bars, 500 µm (a–p); 100 µm (insets in q,r).

**Extended Data Fig. 2:**
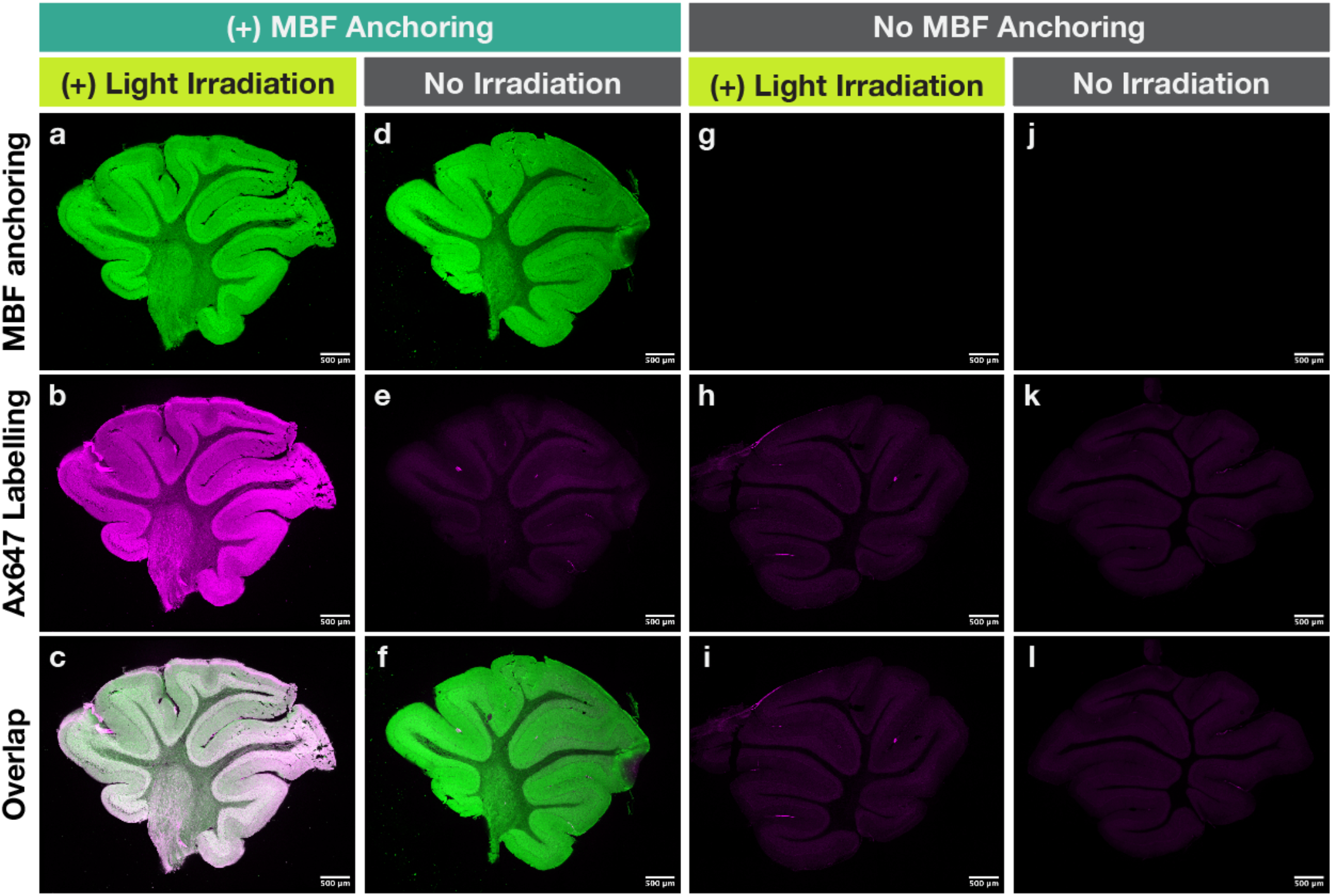
Spatially confined MBF-mediated labelling in cerebellar slices requires both MBF anchoring and light irradiation. **a–c**, Fluorescence imaging of cerebellar slices incubated with MBF–NHS and subjected to DMD-guided light irradiation in the presence of Ax647–hydrazide. MBF anchoring is shown in green (a), Ax647 labelling in magenta (b), and the merged image in c. **d–f**, Control experiment with MBF anchoring but without light irradiation. Minimal Ax647 signal was detected in the absence of irradiation. **g–i**, Control experiment with light irradiation but without MBF anchoring. Negligible Ax647 labelling was observed without the photosensitizer. **j–l**, Negative control without both MBF anchoring and irradiation. No detectable Ax647 signal was observed. These results demonstrate that spatially confined protein labelling in cerebellar tissue requires both MBF anchoring and photoirradiation. Scale bars, 500 µm.

**Extended Data Fig. 3:**
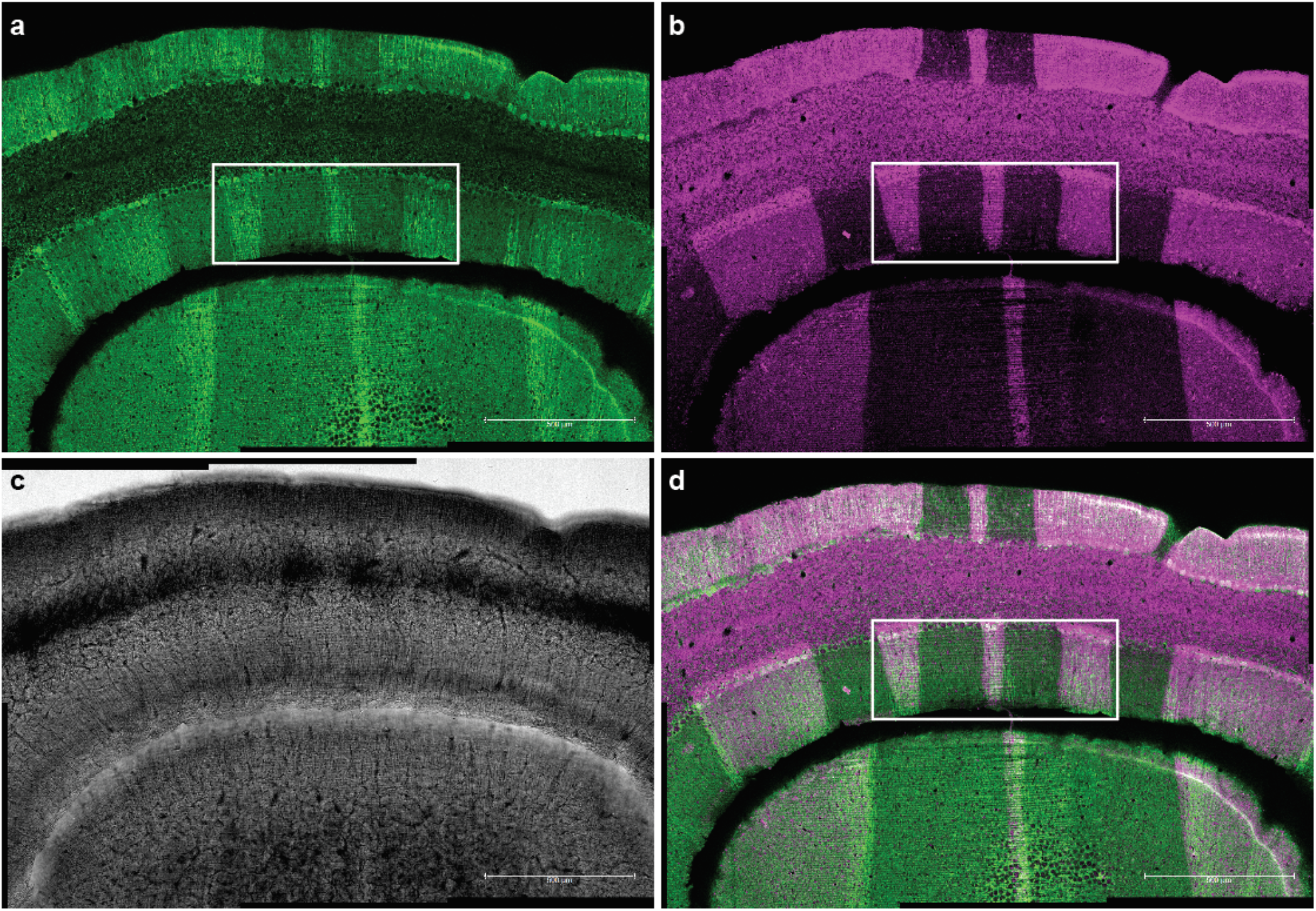
Immunohistochemistry-based ROI definition enables spatially confined labelling in cerebellar slices. **a**, Immunofluorescence image of cerebellar slice showing Zebrin II staining (green), revealing banded anatomical organisation used for ROI definition. **b**, Fluorescence image of Ax647–hydrazide labelling (magenta) following DMD-guided irradiation using ROIs defined from the immunostaining pattern. **c**, Corresponding transmitted light image showing tissue morphology. **d**, Merged image of immunostaining and Ax647 labelling, demonstrating that labelling is confined to the irradiated regions defined by the immunostaining pattern, with minimal signal outside the targeted regions. Boxes indicate regions shown at higher magnification in Fig. 1n–p.

**Extended Data Fig. 4:**
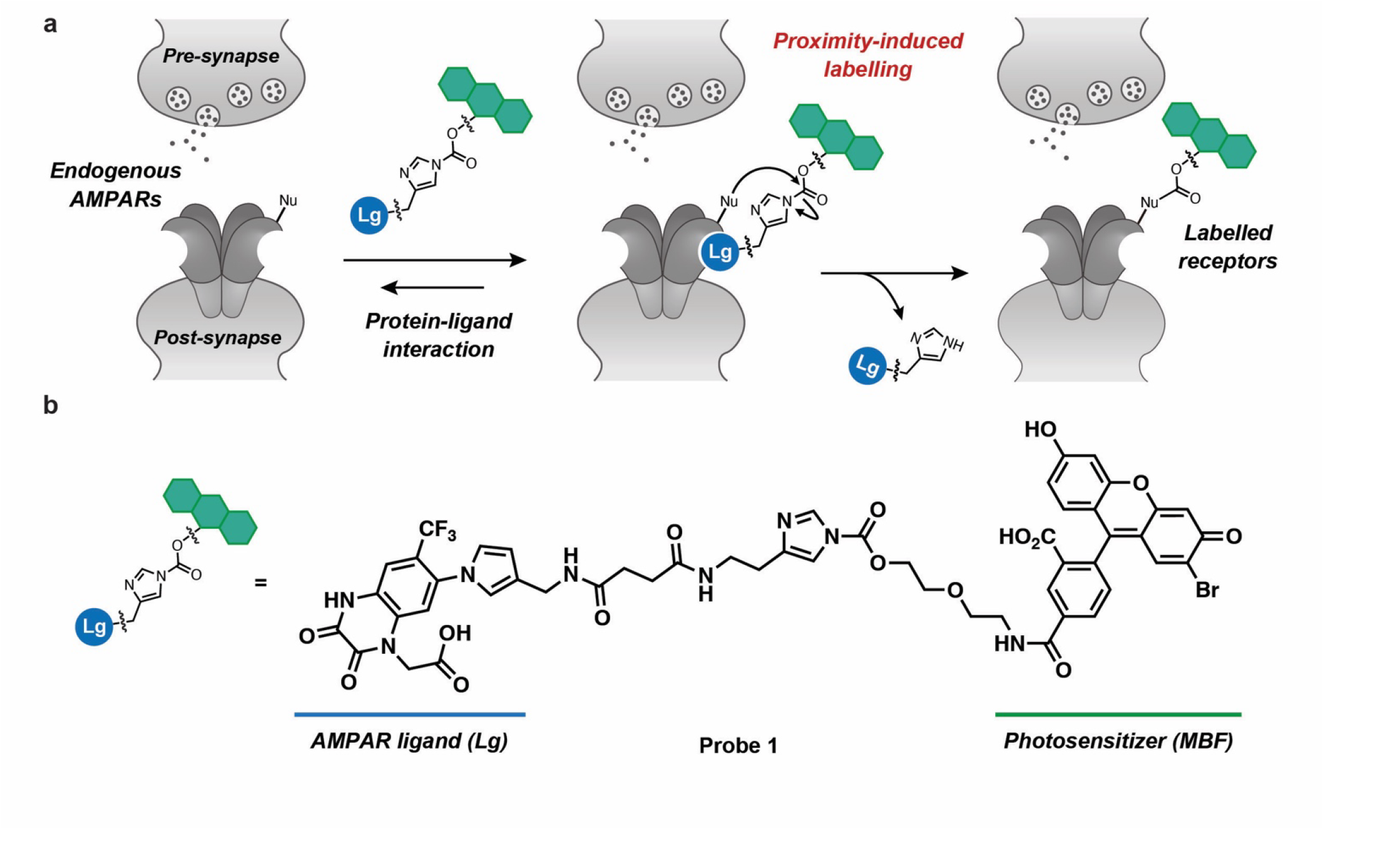
Ligand-directed MBF labelling of endogenous AMPARs for DMD-PhoxID. **a**, Schematic of ligand-directed modification of endogenous AMPA-type glutamate receptors (AMPARs) using an LDAI-based probe. The probe consists of an AMPAR ligand (Lg) linked to a reactive acyl imidazole and the photosensitizer monobromofluorescein (MBF). Upon ligand binding, the probe transfers MBF covalently to nucleophilic residues on AMPARs, enabling protein-centred proximity labelling. **b**, Chemical structure of the LDAI probe (probe 1), comprising the AMPAR ligand, linker, acyl imidazole reactive group, and MBF photosensitizer moiety.

**Extended Data Fig. 5:**
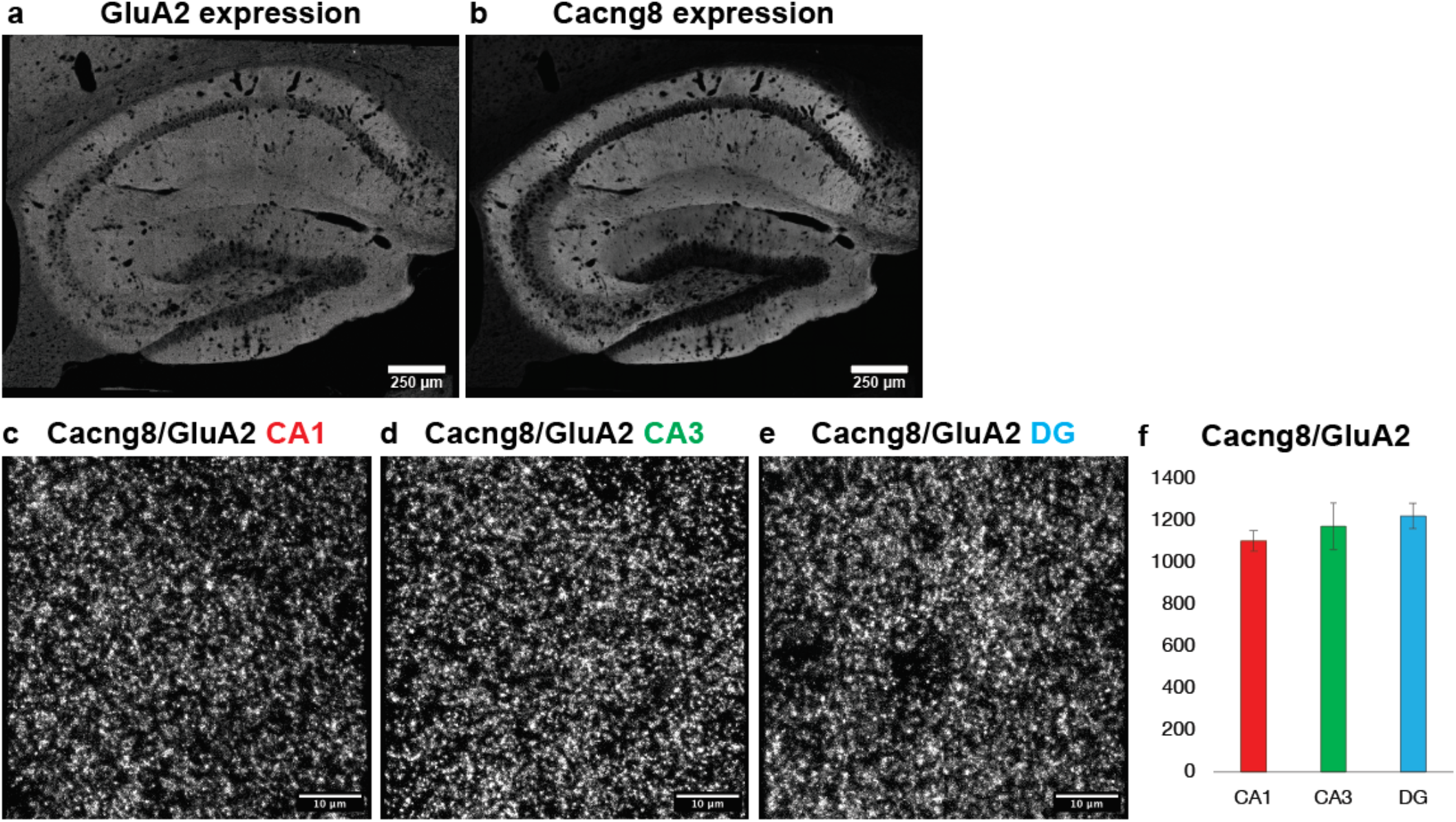
Validation of PLA performance and quantification in hippocampal tissue. **a**,**b**, Immunofluorescence staining of sagittal mouse hippocampal sections showing the distribution of GluA2 (a) and Cacng8 (b) across hippocampal subfields. Both proteins are broadly expressed across CA1, CA3 and dentate gyrus (DG). In addition, Cacng8 is a well-established auxiliary subunit of AMPARs that forms a constitutive complex with GluA2 across excitatory synapses. Together, these features define this pair as a reference expected to exhibit uniform proximity throughout the hippocampus. **c–e**, Proximity ligation assay (PLA) between Cacng8 and GluA2 in CA1 (c), CA3 (d) and DG (e). Representative images show comparable densities of PLA puncta across subfields, consistent with the known constitutive association of Cacng8 with AMPARs. **f**, Quantification of Cacng8–GluA2 PLA signals across hippocampal subfields. Data are presented as mean ± s.e.m., indicating no significant subfield-dependent variation. Together, these results establish that PLA signals can be robustly detected and quantitatively compared across hippocampal subfields under the present experimental conditions, providing a benchmark for interpreting subfield-dependent differences observed for other AMPAR-proximal proteins.

