## Supplementary Information for "Patterned photocatalytic proximity labelling for spatial interactomics"

**Supplementary Table 1: Region-specific proteins identified by spatially confined MBF-based labelling in cerebellar tissue.** Proteins significantly enriched in either the molecular layer (ML) or the granule cell layer (GL) following NHS-MBF-based PhoxID labelling and layer-specific DMD irradiation. "#UP" indicates the number of peptides quantified across replicates. "log2FC" represents the $log_2$-transformed fold change between ML-irradiated and GL-irradiated samples (i.e., ML / GL). Positive values indicate ML enrichment; negative values indicate GL enrichment. Only proteins with ≥2 quantified peptides and consistent enrichment across n = 3 biological replicates are shown.

| Protein | #UP | log2FC | Region | | Protein | #UP | log2FC | Region |
|---|---|---|---|---|---|---|---|---|
| Rbm3 | 2 | -1.175 | GL | | Ctnna3 | 3 | 2.751 | ML |
| Sf3b1 | 6 | -1.065 | GL | | Tf | 2 | 2.541 | ML |
| Smarca2 | 3 | -1.050 | GL | | Prune2 | 2 | 2.190 | ML |
| H2bc12;H2bc14;H2bc18;H2bc3;H2bc4;H2bc7;H2bc9;Hist1h2bp | 6 | -0.991 | GL | | Crkl | 2 | 2.120 | ML |
| Glg1 | 4 | -0.932 | GL | | Hacl1 | 2 | 1.725 | ML |
| Ccsap | 2 | -0.836 | GL | | Chl1 | 4 | 1.470 | ML |
| Hmgb1 | 2 | -0.802 | GL | | Cask | 7 | 1.379 | ML |
| Matr3 | 8 | -0.787 | GL | | Phka1 | 2 | 1.290 | ML |
| Fubp1 | 8 | -0.762 | GL | | L1cam | 6 | 0.937 | ML |
| Trim28 | 4 | -0.731 | GL | | Mpp3 | 3 | 0.815 | ML |
| Hnrnpk | 10 | -0.720 | GL | | Lyn | 4 | 0.754 | ML |
| Ccar2 | 7 | -0.715 | GL | | Shank1 | 16 | 0.731 | ML |
| Gabra6 | 5 | -0.712 | GL | | Erc1 | 6 | 0.720 | ML |
| Ddx17 | 5 | -0.688 | GL | | Cadm3 | 4 | 0.668 | ML |
| Top2b | 5 | -0.686 | GL | | Nrcam | 8 | 0.622 | ML |
| Cars2 | 2 | -0.685 | GL | | Syt12 | 2 | 0.575 | ML |
| Srrm2 | 6 | -0.652 | GL | | Pals2 | 8 | 0.558 | ML |
| Rnmt | 4 | -0.631 | GL | | Nptn | 5 | 0.532 | ML |
| Ik | 2 | -0.627 | GL | | Septin5 | 7 | 0.525 | ML |
| Syne1 | 9 | -0.597 | GL | | Bsn | 25 | 0.507 | ML |
| Dhx9 | 12 | -0.596 | GL | | Cep76 | 3 | 0.502 | ML |
| Sirt2 | 7 | -0.571 | GL | | Wdfy3 | 8 | 0.455 | ML |
| Matr3 | 8 | -0.562 | GL | | Stxbp1 | 24 | 0.454 | ML |
| Nefl | 15 | -0.504 | GL | | Ntm | 4 | 0.433 | ML |
| Gpsm1 | 3 | -0.417 | GL | | Cap2 | 5 | 0.415 | ML |
| Celf2 | 2 | -0.417 | GL | | Tubb4a | 8 | 0.406 | ML |
| Akr7a2 | 4 | -0.411 | GL | | Ank2 | 36 | 0.375 | ML |
| Nif3l1 | 3 | -0.389 | GL | | Llgl1 | 14 | 0.307 | ML |
| Vapb | 6 | -0.376 | GL | | | | | |
| Srpk2 | 4 | -0.362 | GL | | | | | |
| Gclm | 2 | -0.332 | GL | | | | | |
| Ajm1 | 6 | -0.304 | GL | | | | | |
| Sdhb | 4 | -0.237 | GL | | | | | |
| Ckmt1 | 13 | -0.166 | GL | | | | | |
| Fkbp4 | 7 | -0.166 | GL | | | | | |

**Supplementary Table 2: Classification of AMPAR-proximal proteins based on subfield-dependent enrichment, expression patterns and proximity validation.** Proteins annotated to the glutamatergic synapse (GO:0097978) that were enriched by AMPAR-targeted DMD-PhoxID in at least one hippocampal subfield are listed. For each protein, enrichment is quantified as $\log_2$ fold change (log2FC) relative to non-irradiated controls, together with the number of unique peptides detected (#UP) in CA1, CA2/3 and dentate gyrus (DG) samples. Immunostaining patterns across hippocampal subfields are summarized based on previously reported datasets or, where indicated, measurements performed in this study. These data provide an estimate of protein abundance and spatial distribution at the expression level. In contrast, DMD-PhoxID enrichment reflects proximity to AMPARs within each subfield.

| Gene | Protein name | log2FC | | | #UP | | | Immunostaining | | | Ref. |
|---|---|---|---|---|---|---|---|---|---|---|---|
| | | CA1 | CA23 | DG | CA1 | CA23 | DG | CA1 | CA23 | DG | |
| Olfm1 | Noelin | 1.19 | 1.55 | 0.72 | 2 | 2 | 2 | + | + | + | S1 |
| Dlg4 | Disks large homolog 4 | 1.63 | 1.65 | 1.49 | 3 | 2 | 2 | + | + | + | S2 |
| Icam5 | Intercellular adhesion molecule 5 | 2.72 | 2.62 | 2.45 | 5 | 4 | 2 | + | + | + | S1 |
| Tnr | Tenascin-R | 1.22 | 1.13 | 1.07 | 5 | 7 | 4 | + | + | + | S1 |
| Cacng8 | Voltage-dependent calcium channel gamma-8 subunit | 1.54 | 1.44 | 1.23 | 2 | 1 | 1 | + | + | + | S3 |
| Nlgn3 | Neuroligin-3 | 0.92 | 1.38 | 1.19 | 3 | 2 | 2 | + | + | + | S4 |
| Nptx1 | Neuronal pentraxin-1 | 1.08 | 0.90 | | 2 | 2 | | + | + | - | S5 |
| Plxna1 | Plexin-A1 | 1.33 | 0.71 | | 2 | 2 | | + | + | - | Fig.2 |
| Cap1 | Adenylyl cyclase-associated protein 1 | 0.91 | 0.27 | 2.09 | 2 | 2 | 3 | + | + | + | Fig.2 |
| Nrcam | Neuronal cell adhesion molecule | 2.04 | 2.22 | 1.85 | 4 | 5 | 2 | no data | | | - |
| Rims1 | Regulating synaptic membrane exocytosis protein 1 | 1.83 | 0.57 | 0.70 | 2 | 1 | 1 | | | | - |
| Dpysl5 | Dihydropyrimidinase-related protein 5 | 3.06 | 1.98 | 3.58 | 2 | 2 | 2 | | | | - |
| Ywhah | 14-3-3 protein eta | 1.14 | 0.08 | 1.03 | 3 | 2 | 2 | | | | - |
| Shisa7 | Protein shisa-7 | 1.97 | 0.73 | | 2 | 1 | | | | | - |
